# *ImSig*: A resource for the identification and quantification of immune signatures in blood and tissue transcriptomics data

**DOI:** 10.1101/077487

**Authors:** Ajit Johnson Nirmal, Tim Regan, Barbara Bo-Ju Shih, David Arthur Hume, Andrew Harvey Sims, Tom Charles Freeman

## Abstract

The outcome of many diseases is commonly correlated with the immune response at the site of pathology. The ability to monitor the status of the immune system in situ provides a mechanistic understanding of disease progression, a prognostic assessment and a guide for therapeutic intervention. Global transcriptomic data can be deconvoluted to provide an indication of the cell types present and their activation state, but the gene signatures proposed to date are either disease-specific or have been derived from data generated from isolated cell populations. Here we describe an improved set of immune gene signatures, *ImSig*, derived based on their co-expression in blood and tissue datasets. *ImSig* includes validated lists of marker genes for the main immune cell types and a number of core pathways. When used in combination with network analysis, *ImSig* is an accurate and easy to use approach for monitoring immune phenotypes in transcriptomic data derived from clinical samples.

## Introduction

The differentiation and activation of immune cells is associated with changes in the expression of hundreds to thousands of genes (1, 2). Genes specifically expressed by a cell type or cells in a particular state of activation can be used as markers (3) to monitor immune cells in a disease environment, facilitate tailored therapies (4) and clinical stratification of diseases (4, 5). Several studies have identified immune cell markers to ‘deconvolute’ gene expression data. Some of the widely used methods (Table 1) include LLSR (6, 7), qprog (8), DSA (9), PERT (10), MMAD (11) and CIBERSORT (12). For a more detailed review of deconvolution methods read (13). Whilst the derivations differ, all these methods define their marker gene lists based on the comparison of gene expression data from isolated immune cells. Here we employ the principle of co-expression as the basis to derive cell-type specific immune signatures directly from large clinical transcriptomic datasets. This method exploits the fact that the mRNA abundance of genes expressed by a specific cell type will correlate with the number of those cells in a given sample. Between similar samples in a given dataset there are always subtle differences in their cellular composition due to innate variation (e.g. normal variation between individuals, disease severity or subtype etc.), as well as inconsistencies in sampling. Genes expressed by a particular cell type may therefore be identified based on their distinct co-expression profile without the need to physically isolate the cells. We have used this approach to identify robust immune cell type-specific gene expression signatures, known collectively as *ImSig*, derived from and validated on multiple independent datasets to ensure their wide applicability. We have benchmarked *ImSig* against other methods and shown it to out-perform them. We also provide an easy to use algorithm to identify the presence of different immune cell populations in any transcriptomics dataset.

**Table 1.** Summary of the most widely used immune signatures and deconvolution methods

| Authors | Year | Signature derived from | Deconvolution method | No. of Cell types | Total no of markers | Cell types (unique genes) |
| --- | --- | --- | --- | --- | --- | --- |
| Abbas et al. | 2005 | Isolated immune cells | No deconvolution algorithm | 6 | 959 unique genes | B Cell (91), Dendritic Cell (70), Lymphoid (234), Monocyte (82), Myeloid (344), Neutrophil (45), NK Cell (17), T Cell (76) |
| Palmer et al. | 2006 | Isolated immune cells | No deconvolution algorithm | 4 | 1146 unique genes | B cells (427), T cells (241), Granulocytes (411), Lymphocytes (67) |
| Abbas et al. | 2009 | Isolated immune cells | Linear least-squares fits | 17 | 359 Affy u133a probes | Resting helper T cells, Activated helper T cells, Resting cytotoxic T cells, Activated cytotoxic T cells, Resting B cells, Activated B cells, BCR-ligated B cells, IgA/IgG memory B cells, IgM memory B cells, Plasma cells, Resting NK cells, Activated NK cells, Monocytes, Resting dendritic cells, Activated Monocytes, Activated dendritic cells, Neutrophils |
| Nicholas et al. | 2009 | Isolated immune cells | No deconvolution algorithm | 8 | 1842 unique genes | T cells (48), Monocytes (186), B cells (218), NK cells (75), Granulocytes (757), Erythroblast (299), Megakaryocyte (262) |
| Gong et al. | 2011 | Isolated immune cells | Quadratic Programming | Uses signature from other studies |  |  |
| Zhong et al. | 2013 | Isolated immune cells | Linear model & Quadratic Programming | Uses signature from other studies |  |  |
| Newman et al. | 2015 | Isolated immune cells | Support vector machine | 22 | 547 unique genes | B cells naïve, B cells memory, Plasma cells, T cells CD8, T cells CD4 naïve, T cells CD4 memory resting, T cells CD4 memory activated, T cells follicular helper, T cells regulatory (Tregs), T cells gamma delta, NK cells resting, NK cells activated, Monocytes, Macrophages M0, Macrophages M1, Macrophages M2, Dendritic cells resting, Dendritic cells activated, Mast cells resting, Mast cells activated, Neutrophils, Eosinophils |

## Materials and Methods

### Selection of datasets

The primary datasets used for deriving *ImSig* were identified from the Gene Expression Omnibus (National Centre for Biotechnology Information) or ArrayExpress (European Bioinformatics Institute) databases. Datasets from isolated immune cells were identified, and restricted to only those based on the Affymetrix Human Genome U133 Plus 2.0 Array with availability of raw data (.CEL) files. These included: B cells (germinal centre B cells, naïve B cells, memory B cells, IgM+IgD+CD27+ B cells, class switched B cells, IgM+IgD-CD27+ B cells); plasma B cells; monocytes; T cells (central memory T cells, effector memory T cells, naïve T cells, gamma-delta T cells, CD4-T cells, CD4+ T cells, CD8-T cells, CD8+ T cells); macrophages (resting and activated), neutrophils, NK cells and platelets (see Table S5 for details).

A second group of datasets were identified for the purpose of refining and validating the final *ImSig* gene lists. They consisted of blood and tissue datasets, derived from a broad spectrum of diseases and were restricted to data generated on the Affymetrix U133 Plus 2.0 Array with available raw data (see Table S6 for a list of these data).

### Processing of microarray datasets

Quality control (QC) of data from each dataset was performed using the ArrayQualityMetrics package in Bioconductor and scored on the basis of six quality metrics (31). Any array failing more than one metric was removed. Following QC, signal intensity were summarised and normalised using robust multi-array average (RMA) in R using the ‘oligo package’ (32).

Data from isolated immune cell populations were merged and normalised as described above. In order to check that samples clustered according to cell type specific rather than study or any other factor, the RMA normalised data was loaded into the network analysis tool Miru (Kajeka Ltd., Edinburgh, UK). Miru calculates a matrix of pairwise Pearson correlation coefficients (*r*) expression values between every pair of genes/samples in a dataset. Graph layout in-tool is performed using a modified Fast Multipole Multilevel Method (FMMM) (33) and the resulting network is rendered in a 3-D environment. Networks are composed of nodes (representing transcripts/samples) connected by weighted edges (representing correlation values). After loading the immune cell data a sample similarity network was then plotted at a correlation threshold of *r* > 0.83. All sample outliers i.e. samples that did not group with other samples of the sample type, were removed. The remaining 329 samples clustered based on cell type rather than study (Figure S4). For blood and tissue datasets, the data was collapsed to one probe-set per gene by choosing the probe-set with the highest variance across samples.

### Refinement of signatures (Cluster model algorithm)

The quality-controlled datasets were loaded into the network analysis tool Miru. Within the tool, a correlation network was generated and clustered using the MCL algorithm (inflation value: 2.2). A proportion of the genes in each MCL cluster were replaced with random genes in increments of 2% from 0-100% of total genes. The percentage of genes from the original MCL cluster in this modified cluster was defined as Percent_similar_ (percentage of genes with high similarity). The similarity of each gene to other members of the cluster or annotation is defined by the median value of its Pearson correlation coefficients to every other member within a cluster or annotation, and the median of this value from all genes within a cluster or annotation is referred to as Pearson_group_. The decrease in Pearson_group_ with increasing replacements of MCL cluster by random genes was modelled as a sigmoid function of Percent_similar_ using nonlinear least squares in R.

In the situation where the groups of genes were of the same signature (different cell type signatures derived from both blood and tissue) instead of modified MCL clusters, the Percent_similar_ is unknown whilst Pearson_group_ can be calculated. Therefore, inverse estimates of Percent_similar_ using Pearson_group_ were made using the R package “investr”. The upper and lower threshold of Pearson_group_, beyond which investr function cannot estimate the Percent_similar_, were noted and used as cut off for determining if genes will be discarded from the refined signature.

In the second stage of the filtering process, signatures with a Pearson_group_ 1) higher than upper threshold were left unchanged; 2) between upper and lower thresholds were reduced in size, using the model above to determine the number of genes to discard; 3) less than the lower threshold were considered to be absent from the dataset. This method of filtering would allow greater stability cross datasets, whilst retaining more flexibility with a more comprehensive list of genes with informative signature. We used the cluster model algorithm on eight blood and eight tissue datasets to refine the *ImSig* signature lists (Figure 2A)

**Figure 2:**
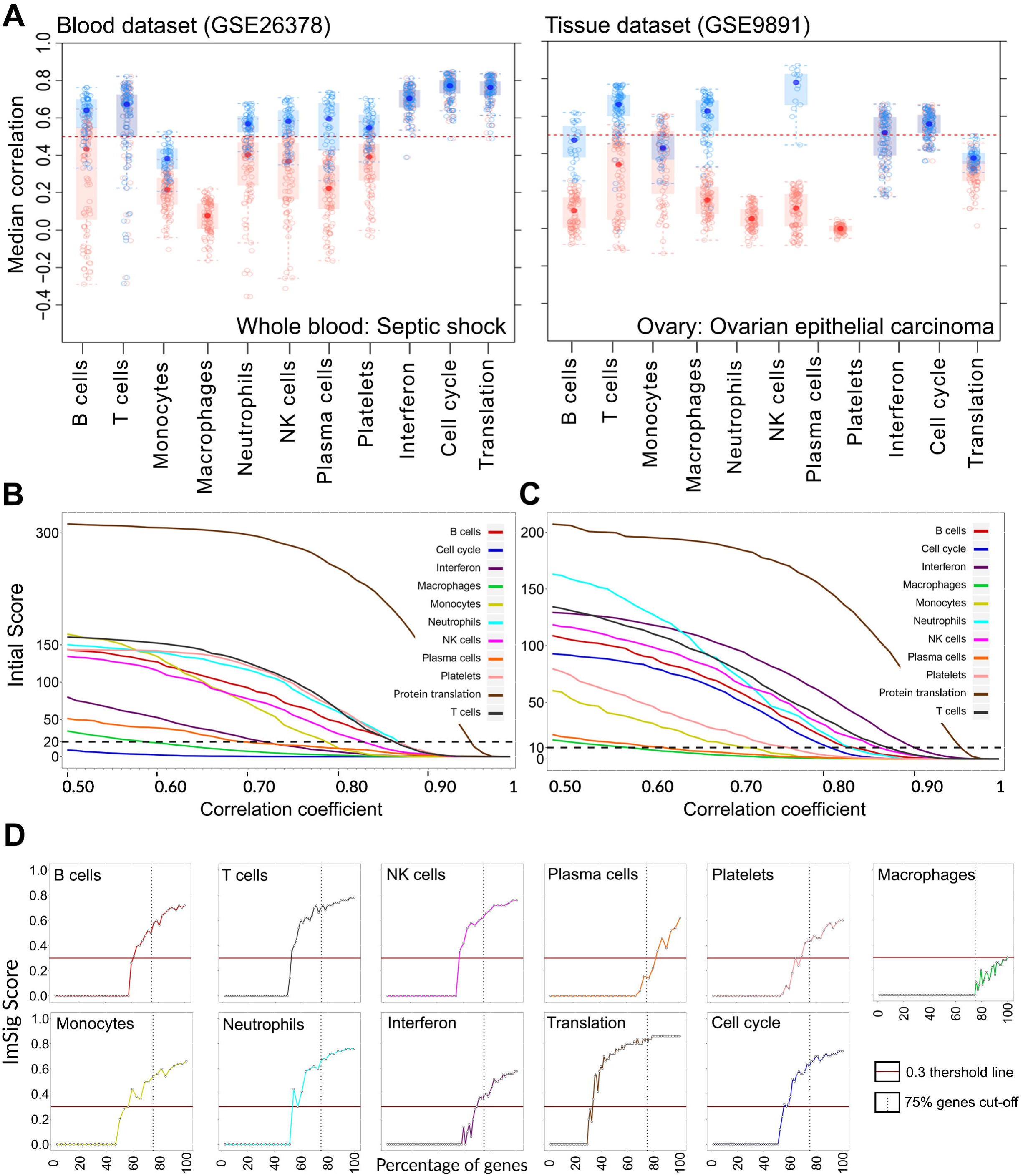
Cluster model algorithm refinement and *ImSig* algorithm. **A)** The plots represents the outcome of running the cluster model algorithm over a blood and a tissue dataset. Each node represents a unique gene and plotted as a function of its median correlation value within the signature. Blue colour represents the genes that were kept and red represents the genes that were discarded after running the algorithm. The algorithm was applied to eight blood and eight tissue datasets (only 2 shown above). All the blue nodes were then pooled to identify the most commonly occurring genes across datasets, which then formed the basis of defining *ImSig*. **B and C**, Line plots showing‘initial score’ calculated for every correlation cut-off between 0.50 and 0.99 while calculating the *ImSig* score. For **B)** microarray dataset (heart attack, GSE48060), the threshold line is drawn at 20 and for **C)** RNA-seq dataset (Brucellosis; E-GEOD-69597), the threshold line is drawn at 10. **D)** Plots showing the effect of loss of signature genes on *ImSig* score. These were calculated by performing a permutation analysis of removing signature genes randomly.

### Derivation of ImSig_blood_

The most differentially expressed genes (DEGs) for each isolated immune cell type was determined by calculating the average fold change for a particular cell type relative to the rest. The top 100 DEGs for each cell type were refined across eight blood datasets (Table S6) using the cluster model algorithm. The resultant sets of genes derived from each dataset were then compared, and the most overlapping set of genes were defined as the blood signature set, *ImSig*_blood_.

### Derivation of ImSig_tissue_

The same approach as of *ImSig*_blood_ was not successful in defining a tissue specific *ImSig*, since the top 100 DEGs for each cell type were poorly co-expressed in complex tissue datasets. This is likely due to the fact that these 100 DEGs were derived from isolated cells, some of which were cultured *in vitro*, where their phenotype more closely resembled that their counterparts in blood. We therefore decided that the best approach was to use a correlation based approach. The expression data of isolated immune cells was loaded into the network analysis tool Miru. A large and highly structured network graph was constructed using a correlation threshold of *r* > 0.8. The network was then clustered into groups of genes sharing similar profiles using the Markov Clustering (MCL) algorithm with an MCL inflation value set to 2.2 (34). These clusters were then extensively explored to find genes that were distinctively expressed in only one cell type in contrast to the rest. These genes were then explored in the context of four tissue datasets as a class set and network graphs constructed and clustered as described earlier. For each dataset, clusters identified as being specific (based on the added class set) to a particular cell type were isolated. The resultant set of genes were compared to each other and the most common set of genes were refined in another 8 tissue datasets (Table S6) using the cluster model algorithm to define the *ImSig*_tissue_.

### Derivation of pathway signatures

Whilst analysing the clusters and refining them to be cell type-specific, we also identified a number of other clusters that were consistently co-expressed across different datasets. With the help of GO Annotation and known marker genes, we were also able to define these clusters as cell cycle-associated, interferon stimulated and protein translational activity. These clusters were further refined in blood and tissue datasets as describe above using the cluster model algorithm.

### Validation of ImSig in mixed cell population datasets

*ImSig* was validated using additional independent datasets, including two blood (heart attack blood samples: GSE48060 and type I diabetes mellitus blood samples: GSE55098), two tissue (breast tumour tissue samples: GSE58812 and primary CNS tumour tissue samples) and an infection dataset (*Chlamydia trachomatis* infection tissue sample: GSE20436). All datasets were pre-processed as described above. A number of transcriptomic profiles derived from RNA-seq technology were also analysed by *ImSig* to ensure its wide applicability and lack of platform dependency, in particular RNA-seq data were downloaded from TCGA database.

### ImSig cluster scoring algorithm

In order to facilitate the use of *ImSig* a scoring system was devised that supports the identification of any given signature without the need to perform network analysis. For any given transcriptomic dataset, the calculation of the *ImSig* scores is a two-step process where an initial score is first computed based on the following formula:

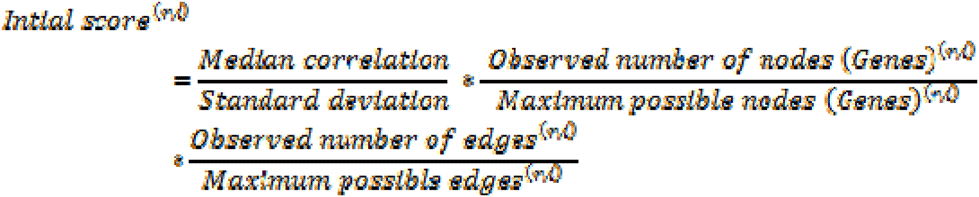

Where *r* is the correlation cut-off and *i* is the cell type/pathway signature.

Median correlation is calculated by computing the correlation values across samples for all possible pairs of genes within any given signature and then taking the median value. The standard deviation is calculated by computing the mean expression value of all genes within a signature and then calculating its standard deviation across samples. The maximum possible edges is calculated with [n*(n-1)]/2, where *n* is the number of genes in any given signature. The maximum possible nodes is the number of genes defining a particular *ImSig* signature. The initial score is computed for all eight cell type clusters (B cells, T cells, monocytes, macrophages, NK cells, neutrophils, plasma cells, platelets) and three pathway clusters (cell division, protein translational and interferon response) using a range of Pearson correlation coefficient thresholds, from 0.50 to 0.99 at 0.01 intervals. The resulting matrix contains 50 scores for each of the signature. At this point we set an ‘initial score threshold’ of 20 and 10 for microarray and RNA-seq datasets, respectively (these were determined empirically, Figure 2B&C). Any value below this threshold is not regarded to be a genuine cluster due to a poor correlation between genes within the signature at the set *r*-value. We recommend these thresholds as they are based on observations from numerous datasets. Following this the final *ImSig* score is calculated for each cluster using the following formula.

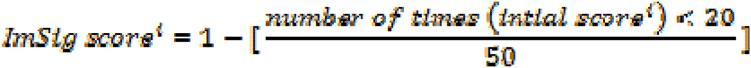

All data should be in log scale for calculating *ImSig* score. An R script is available for running *ImSig* scoring algorithm. The script can be downloaded here: www.github.com/systems-immunology-roslin-institute/ImSig. The final score (*ImSig* score) is a value between 0 and 1. After extensive evaluation, any value above 0.3 is regarded as evidence that the cell type/pathway signature is present in the dataset.

### Comparison with CIBERSORT

A blood and a tissue dataset were used for this purpose where there was some prior knowledge about the cell populations present and their relative abundance in sample sub-groups.

A blood dataset (SLE patients: GSE49454) was downloaded from GEO. The authors of this study had provided the cell counts along with the transcriptomics data in this file. Initially, the patients were ordered based on the cell count for each of the different cell types independently (B cells, T cells, NK cells and neutrophils). They were then equally divided into three groups and the top and bottom groups were used for analysis. Two-tailed unequal variance T-test showed a significant alteration in cell counts between these two group of patients for all four cell types (p<0.05). Using CIBERSORT and *ImSig* the relative proportion of immune cells were then computed. For CIBERSORT the data was loaded into (https://cibersort.stanford.edu/) as per the authors instructions and the computed relative proportions were downloaded. The relative proportions of immune subtypes were all summed to make up the parent cell type (T cells, B cells, neutrophils, NK cells). Then, each cell type was normalised independently to be represented as a fraction of 1 across samples (i.e., the sum of normalised cell proportion for any cell type is equal to 1). Similarly, for *ImSig* the relative abundance of immune cells were calculated by averaging the expression of signature genes for each sample and then normalised to represent them as a fraction of 1. Two-tailed unequal variance T-test was then used to test for significant change in cell proportions between the two groups of patients in all four cell types.

Similarly, a tissue dataset (trachoma: GSE20436) was downloaded. The patients were divided into three groups as per the level of infectivity according to its authors (controls, symptom +ve/*C. trachomatis* −ve patients, and symptom +ve/*C. trachomatis* +ve patients). As described earlier, the relative proportion of immune cells (T cells, B cells, neutrophils, monocytes, macrophages, NK cells and plasma cells) were computed and normalised using CIBERSORT and *ImSig*. This was followed by a one-way analysis of variance (ANOVA) to test for significant changes in cell numbers between the three groups of patients.

## Results

### Blood and tissue immune signatures (ImSig_blood/tissue_)

*ImSig* was derived as described in the experimental procedures, and as shown in (Figure 1). Briefly, an initial meta-analysis was carried out on 330 samples of isolated human immune cell populations and the top 100 differentially expressed genes were determined for each immune cell type. Using a network-based approach to identify sets of robustly co-expressed (correlated) genes in a variety of blood datasets, the lists were further refined (Figure 2A). The resulting cell-specific marker gene lists were collectively named *ImSig*_blood_. However, the limitations of this approach become evident from network analysis of clinical tissue transcriptomic datasets, where the cell-based marker genes showed independent expression. To overcome this issue we identified the most conserved cell type-specific groups of genes based on their co-expression across four tissue datasets, and further refined them by examining a eight other tissue datasets (Figure 2A). This resulted in our *ImSig*_tissue_ gene signatures. *ImSig*_blood_ contains 491 marker genes and *ImSig*_tissue_ contains 569 marker genes for B cells, monocytes, macrophages (tissue only), neutrophils, NK cells, T cells, plasma cells, platelets (blood only), cell cycle, protein translation and interferon signalling. For a full list of the genes comprising these signatures and numbers for each cell type or pathway see Table S1. GO term analysis confirmed that the cell marker lists for both *ImSig* signatures were highly enriched in genes related to immune function (Table S2, S3). The overlap between blood and tissue signature varied depending on cell type/pathway (Figure S3).

**Figure 1:**
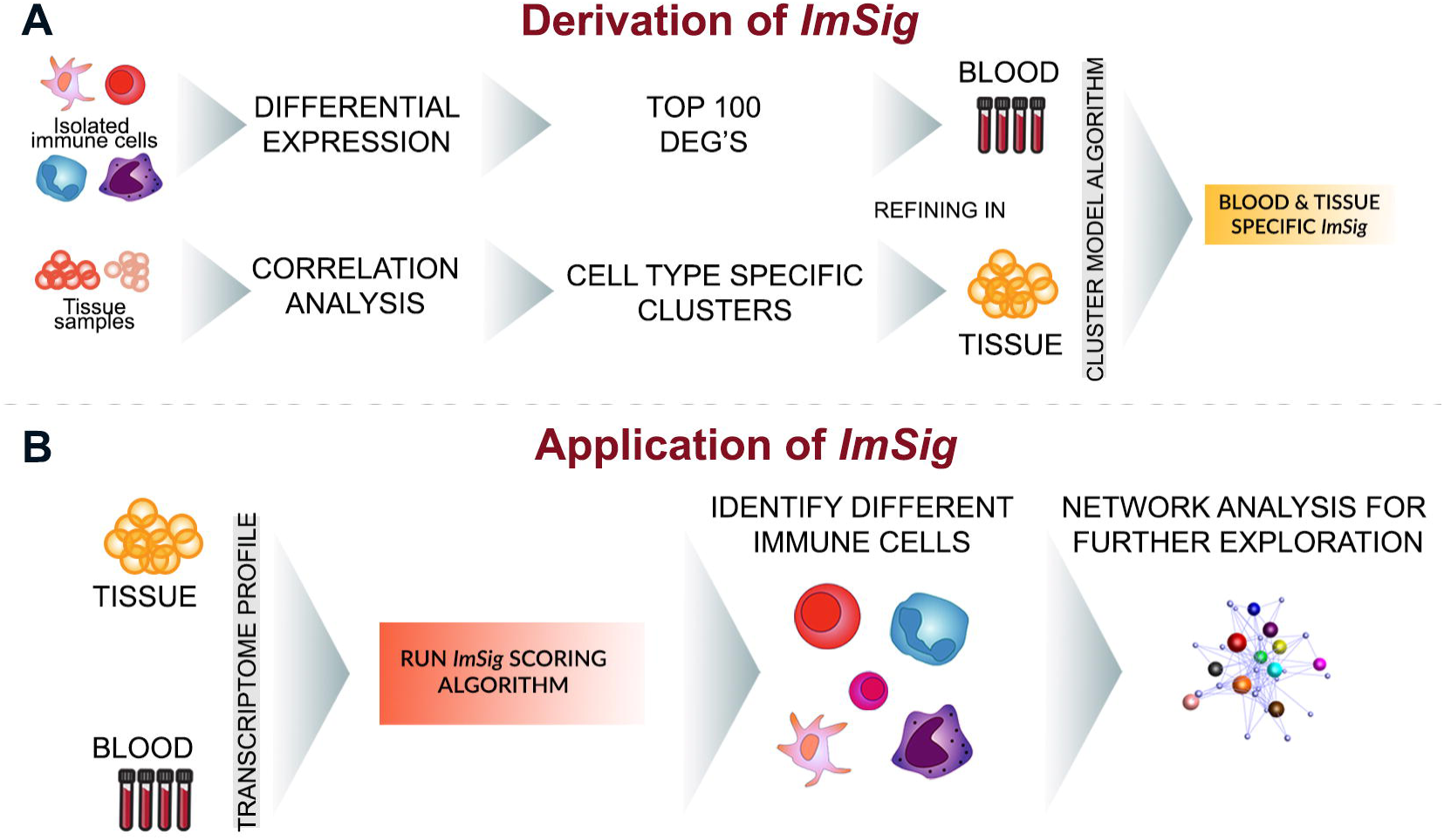
Derivation and application of blood and tissue *ImSig*. **A)** Flow chart depicts the systematic derivation of *ImSig*. The transcriptome of isolated immune cells was subjected to differential gene expression analysis or correlation analysis to derive a preliminary list. This was further refined using the cluster model algorithm to define the blood and tissue-specific immune signatures (*ImSig*). **B)** Application of signatures involves running *ImSig* scoring algorithm on any transcriptomic data to identify the different immune cells present within the samples followed by network analysis to study the genes that are correlated best with the core signature genes.

### Genes that make up the signatures

Table S1 highlights the sets of genes that distinguish *ImSig*_blood_ and *ImSig*_tissue_. The process signatures; cell cycle, interferon response and protein synthesis (translational activity) are relatively robust in both blood and tissue. The T cell clusters in both cases are anchored and validated by the subunits of CD3, but otherwise, there is very little overlap. The implication is that the T cells that enter tissues in a pathological situation are radically different in their gene expression profiles from the bulk of naïve T cells in peripheral blood. Note that there is no evidence of a cluster of genes associated with specific T cell polarisation states. The key transcription factors *FOXP3* (Treg), *RORC* (Th17) and *GATA3* (Th2) do not form part of clusters, since they are expressed by other cell types. However T-BET (*TBX21*), considered to be a Th1 specific transcription factor is in the NK cell cluster a cell type in which it is also strongly expressed. The NK cell cluster also shows considerable divergence between blood and tissue, in particular the NK cell receptor family being much more robustly co-expressed in tissue RNA. In blood, many of these receptors are also detectable in gamma-delta T cells (14). The various myeloid clusters are rather more difficult to be associated with specific cell types. The macrophage cluster, specific to the tissue data set, contains the CSF1R, which is known to be macrophage-specific and essential for differentiation and survival (15), and also contains many of the genes that are up-regulated in monocyte-derived macrophages derived by cultivation in CSF1 (16). An unexpected member of this cluster is CD4. In blood, CD4 is expressed at similar levels in CD4+ T cells and monocytes, and so does not form part of a T cell cluster. In tissue, CD4 is highly-expressed by macrophages, and correlates more highly with their presence than with the presence of T cells. The clusters annotated provisionally as monocyte and neutrophil have very little overlap between the blood and tissue profiles. Archetypal markers, CD14 for monocytes and the G-CSF receptor and chemokine receptor CXCR2 (the receptor for IL8) are co-expressed with very different gene sets in blood and tissues. Hence, it may be more appropriate to consider distinct separate myelomonocytic regulons, reflecting the rapid differentiation of these cells following extravasation. For example, S100A8/A9, which encode the most abundant neutrophil proteins (17), are also expressed by monocytes, but rapidly down-regulated as they differentiate to macrophages. The mRNAs encoding many neutrophil-specific granule proteins (MPO, lactoferrin etc) are expressed most highly in progenitor cells (16), and do not contribute to a signature in either blood or tissue.

### ImSig scoring algorithm

The *ImSig* scoring algorithm was developed to reflect the correlation and expression level of the marker genes in any given dataset. The algorithm generates a numerical likelihood score that a given signature is present in a dataset. Based upon empirical evaluation of a wide range of data, an *ImSig* score >0.3 indicates positive identification of the signature in a given dataset (Figure 2B&C). *ImSig* scores for all the validation datasets along with six other RNA-seq datasets are provided in Table S4. Consistent with its derivation, the *ImSig* macrophage signature is absent from any blood datasets, irrespective of platform. Conversely, the platelet signature was not scored positive in any tissue dataset examined. As with other deconvolution methods, *ImSig* works best when the majority of signature genes are present in the dataset to be analysed. Based upon a permutation analysis of the effect of random removal of genes on the *ImSig* score (Figure 2D) a minimum of 75% of the genes from each individual signatures is required for an accurate representation analysis. Being correlation-based, a dataset generally needs to comprise of at least 20 distinct samples is needed to provide sufficient diversity before the *ImSig* algorithm can be applied.

### Validation of blood and tissue marker genes

To test its universality, we applied *ImSig*_blood_ to deconvolution of a range transcriptomics data derived from whole blood or peripheral blood mononuclear cells (PBMC). Examples of these analyses are given here. Data from the blood of 21 control and 31 heart attack patients (GSE48060) identified the presence of B cells, T cells, NK cells, plasma cells, platelets, monocytes and neutrophils (Figure S1A). In terms of the average expression of marker genes, no consistent difference was observed between the control and heart attack samples suggesting that relative blood cell numbers were not altered. The macrophage and cell cycle signatures were not detected. In contrast, *ImSig* analysis of PBMC’s from control and patients with type 1 diabetes mellitus (GSE55098) identified increased proliferation in a number of samples (Figure 3A) and the analysis also clearly identified the presence of T cells, B cells, along with plasma cells, monocytes, neutrophils, NK cells and platelets (Table S4). Notably there was also significantly lower expression (p=1E-10) of the NK cells markers genes in samples derived type 1 diabetes (Figure 3A) where these cells are known to be dysregulated (18, 19).

**Figure 3:**
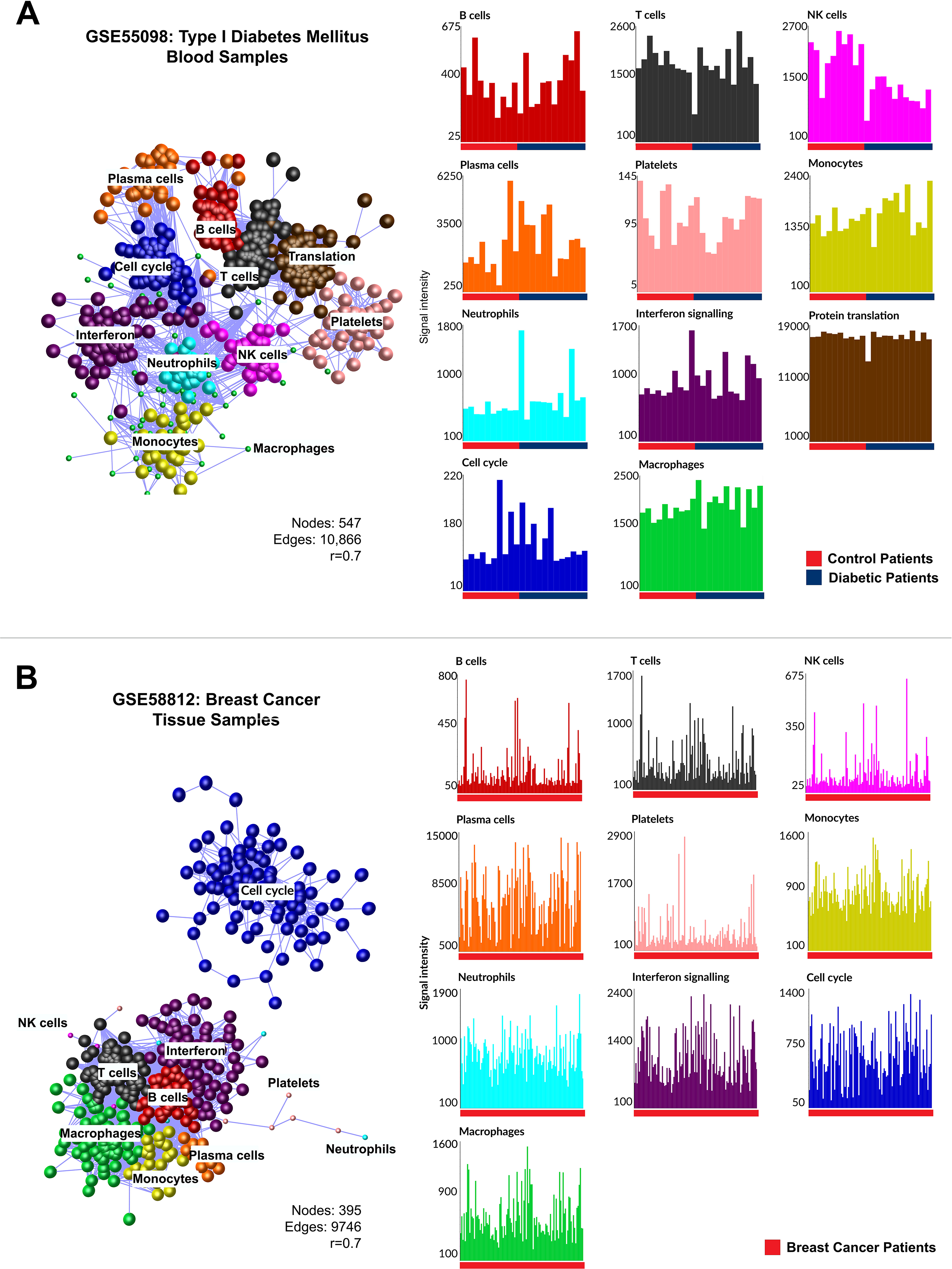
Deconvolution of blood and tissue datasets. **A)** Correlation network of gene expression data from blood samples of patients with type I diabetes mellitus represented and **B)** samples from breast cancer patients. Each cluster represents a unique cell type. Nodes derived from other signatures which were included in the graph but did not cluster are reduced in size. Histogram plots represent the average expression profile of the *ImSig* signatures across samples.

To validate *ImSig*_tissue_, we first examined a dataset of triple-negative breast cancers derived from 107 patients (GSE58812). As expected, and in keeping with our previous network analysis of multiple tumour datasets (20), the cell cycle cluster was readily detected, reflecting the heterogeneity in proliferative index between tumours. The analysis revealed macrophages, T cells, B cells, plasma cells, interferon but there was no evidence of platelets, neutrophils and NK cells present in these samples (Figure 3B). The levels of all immune cells (as judged by the average expression of the marker genes) varied greatly between samples. By contrast, a relatively small brain tumour dataset comprising 23 samples of primitive neuroectodermal tumors and medulloblastomas lacked evidence of immune cell infiltration, other than an NK signature (Figure S1B). Being behind the blood-brain barrier, lymphocyte populations in these tumours are likely absent or at very low levels (21) but infiltration of T cells was evident in other brain tumour datasets that we have analysed (Table S4). Neutrophil signatures were absent from tumour datasets. However, as expected, a dataset of eye swabs taken from eyes of controls or children with the symptoms of trachoma (GSE20436) (22) was positive for all signatures of immune cells (Figure S2). Previous studies have shown that in certain chlamydial infections, neutrophils recruit T cells to the site of infection (23), other studies report the involvement of NK cells, monocytes and macrophages (24–26). Finally, we demonstrate the explorative power of *ImSig* when coupled with network analysis. The genes comprising the signatures were selected as being core ‘invariant’ markers of a particular cell type. When used in the context of a correlation analysis of a complete dataset, if the relevant cells are present within the samples, surrounding the signature genes will be other genes expressed in these populations. In this manner one can better evaluate the activation state of immune cells *in situ*. Using the trachoma dataset as an example we highlight known immune related genes that were co-expressed with *ImSig* core signature genes (Figure 4). The associated *ImSig* scores for all the validation datasets can be found in Table S4.

**Figure 4:**
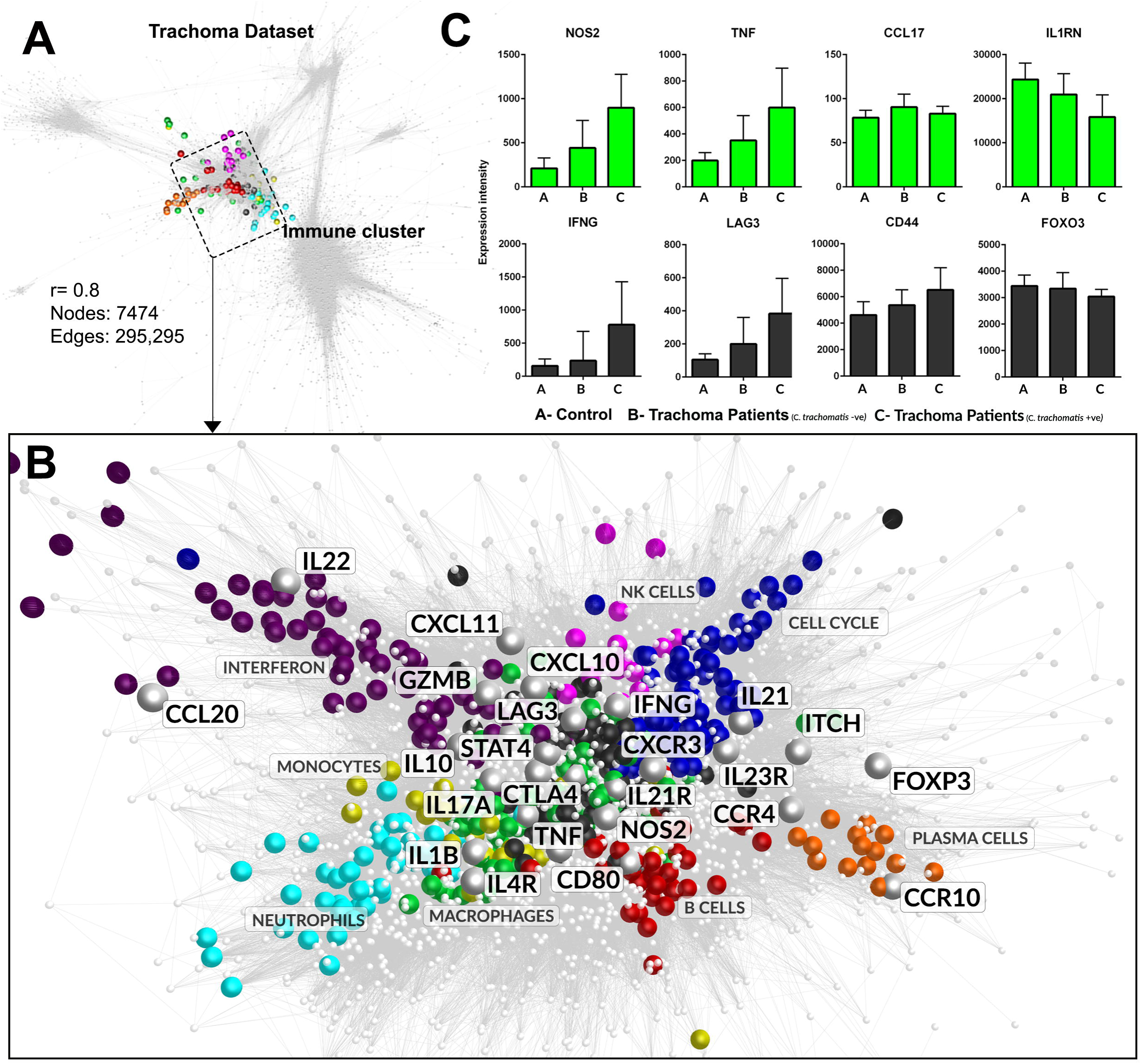
Network graph to highlight a few closely correlated immune related genes with *ImSig*. **A)** Correlation network of gene expression data from trachomatis infection (GSE20436). The nodes represent unique genes and the *ImSig* genes are coloured to highlight the immune cluster. **B)** A close up of the immune cluster. The *ImSig* related genes are coloured to represent different immune cell types, while the remaining genes are reduced in node size. We highlight a few well known immune modulatory genes with a greater node size and marking their gene symbols alongside. **C)** Bar plots represents the average expression intensity of individual genes across samples. The top panel (Green) plots represents a few marker genes to understand macrophage biology and the bottom panel (dark grey) to understand the T cell biology.

### Comparison with CIBERSORT

The ability of *ImSig* and CIBERSORT to identify changes in relative proportions of cells between sample groups was compared using a blood (GSE49454: Systemic lupus erythematosus patients) and a tissue dataset (GSE20436: trachoma). For the blood dataset, cell counts were available for B cells, neutrophils, T cells and NK cells. Both methods generally performed well, *ImSig*_blood_ demonstrated a significant difference (p<0.05) in all four cell types, although CIBERSORT failed to show a significant difference in B cells (p=0.389) (Figure 5A, Table S7). Samples from the trachoma dataset were divided into three groups of 20, based on the level of infection as originally described (for more detail see Methods). Although actual cell counts are not available for these data, it is known that the immune infiltrate increases with the level of infection (27). *ImSig*_tissue_ showed there to be a significant increase (p<0.05) in all seven cell immune types (B cells, neutrophils, T cells, NK cells, plasma cells, monocytes and macrophages) during an active infection, while significant differences were only reported for T cells and macrophages using CIBERSORT (Figure 5B, Table S7). Moreover, the pattern observed using CIBERSORT did not seem to correlate with the infection status of C. *trachomatis* (Figure 5B). CIBERSORT was also used in its native form, i.e. the subtypes were not summed to represent the parent population. A significant change in cell number was observed only for M2 macrophages (p=0.001), activated mast cells (p=0.022) and resting dendritic cells (p=0.0007). The 19 other immune cell groups defined by CIBERSORT showed no significant difference in cell proportion across patient groups (Table S8).

**Figure 5:**
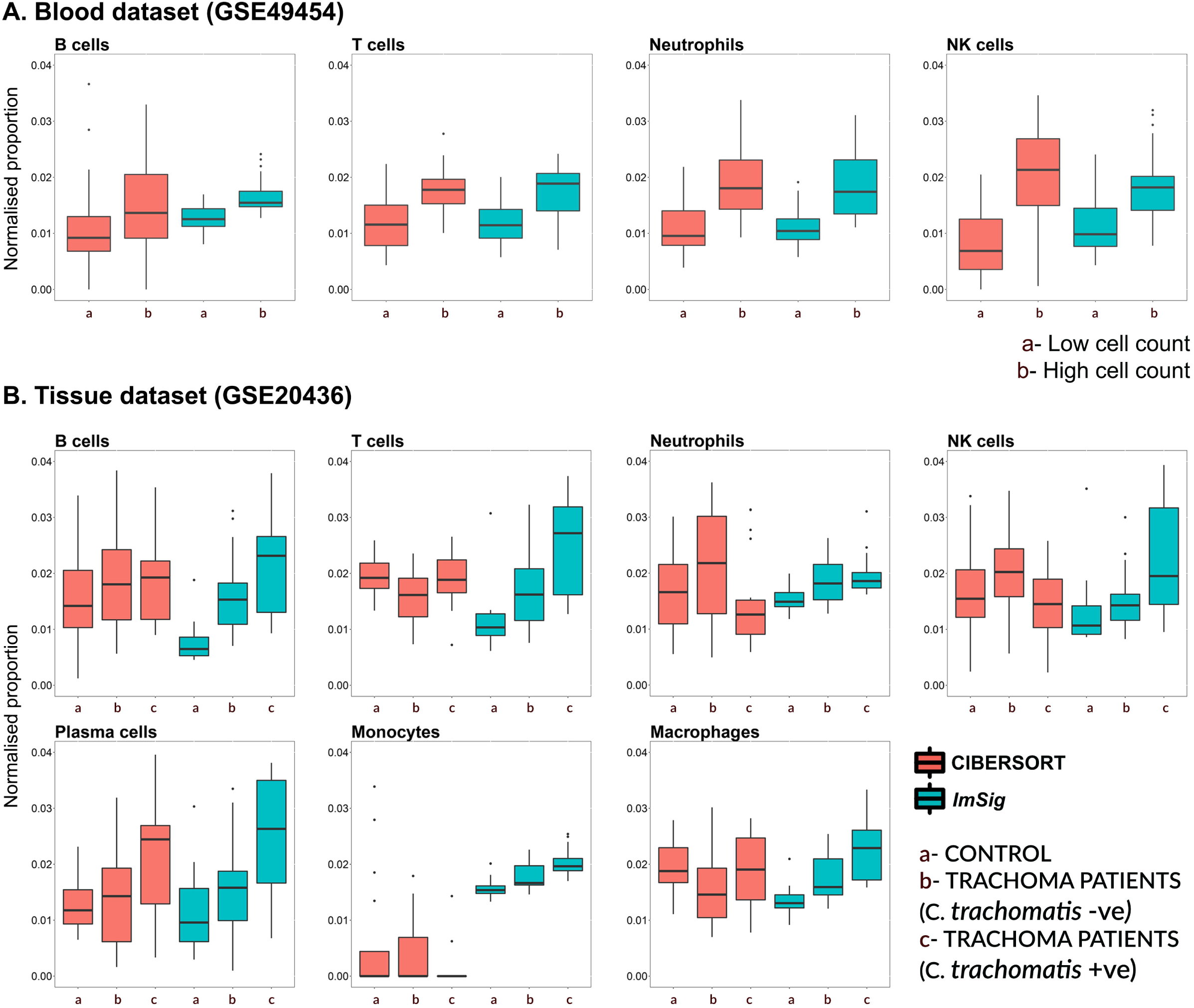
Comparison of *ImSig* with CIBERSORT. **A)** Comparison performed using a blood dataset. The boxplots show the relative abundance of immune in cells in the two patient groups computed by CIBERSORT and *ImSig*. The actual median cell count for the four immune cell types were (high, low) Neutrophils (2655, 6160), T cells (617.5, 1988), B cells (35, 293) & NK cells (22.5, 176.5). Significant difference was observed for T cells, Neutrophils and NK cells using CIBERSORT while all differences seen in *ImSig* including B cells are significant (P value <0.05). **B)** Comparison performed using a tissue dataset. The boxplots show the relative abundance of immune in cells in the three different patient groups computed by CIBERSORT and *ImSig*. Significant difference was observed only for macrophages and T cells using CIBERSORT while all differences seen in *ImSig* are significant (P value <0.05).

## Discussion

In the last few years a number of immune marker gene signatures have been proposed (6-12). The current work is based on the observation that when correlation (co-expression) network analysis is employed to explore large transcriptomics datasets derived from normal or diseased tissues, clusters of genes associated with specific immune cell populations, or specific transcriptional regulons such as protein synthesis, interferon response or cell cycle, are frequently observed clustered together (20, 22, 28, 29). This is because the abundance of mRNAs derived from cell-specific, or process-specific genes is correlated with relative number of those cells expressing those genes within a sample, resulting in their observed co-expression across a sample set. The most important conclusion from our analysis is that signatures based upon cells isolated from blood cannot be applied with any confidence to tissue data.

The utility of the blood and tissue *ImSig* gene lists has been demonstrated through applications to a number of datasets. Other approaches to deconvolution include LLSR (7), qprog (8), DSA (9), PERT (10), MMAD (11) and CIBERSORT (12). Each is based on a signature derived by a different data mining approach ranging from simple matrix decomposition to complex iterative procedure. Of these methods CIBERSORT was shown to out-perform others (12) in terms of analysis of tissue data with noise or unknown content and was reported to be able to differentiate closely related cell types. CIBERSORT includes profiles for 22 distinct cell types, including various states of T cell activation and macrophage differentiation. The network analysis of disease datasets herein does not support robust clusters that distinguish macrophage activation states, in keeping with previous analysis (20). In essence, the best one can do is define three myeloid states (neutrophil, monocyte, macrophage), and the inducible genes are disease/lesion specific. Expression QTL analysis of inducible gene expression in monocytes suggests that inducible gene expression profiles may also be individual-specific (30).

An ideal workflow for employing *ImSig* would involve running the *ImSig* algorithm to identify the different immune cell populations in a dataset and then using the average expression of signature genes to understand the relative proportion of cells between samples and clinical subsets. This can be followed by network analysis which can be used to better understand the wider context of the immune environment. Through observing the genes that closely correlate with the core signature genes, one can better under the type of activation or indeed the level of involvement which these cells play in a given microenvironment of a disease state. As an example we have highlighted a few immune related genes that are co-expressed with our core signature genes in the trachoma dataset (Figure 4). The expression profiles of known immune modulatory genes such as *IFNG, LAG3, CD44, FOX03, FOXP3, CD80, IL20, STAT4, IL17A* etc are correlated with the core macrophage and T cell signature genes, suggesting that the macrophages are undergoing classical activation, and the T cells include Th17, TReg and Th1 states. Thus such explorative analysis can be employed using *ImSig* to understand the differentiation state of immune cells between patient groups.

The *ImSig* algorithm has been tested on data derived microarray and RNA-seq platforms. We have also tested its applicability across a wide range of datasets derived from blood, tissue, sputum and faecal samples (data not shown). As long as immune cells are present, *ImSig* efficiently identifies the cell types present. We therefore anticipate that *ImSig* and the methodological approaches described here will prove valuable for studying immune cell variation in human transcriptomics data derived from a wide variety of conditions clinical samples.

## Author contributions

A.J.N performed the majority of work described here. A.J.N, T.R, B.J.S, D.A.H, A.H.S and T.C.F wrote and edited the manuscript. A.H.S and T.C.F supervised the project.

## Acknowledgements

A.J.N is a recipient of The Roslin Institute and CMVM scholarship and Edinburgh Global Research Scholarship. A.H.S is grateful for funding from Breast Cancer Now. T.R, B.J.S and T.C.F are funded by MRC consortium grants (MR/M003833/1, MR/L014815/1) and T.C.F is funded by an Institute Strategic Grant from the Biotechnology and Biological Sciences Research Council (BBSRC) (BB/JO1446X/1). The authors have no conflict of interest.

